# Next-generation biocomputing: mimicking artificial neural network with genetic circuits

**DOI:** 10.1101/2021.03.12.435120

**Authors:** Leo Chi U Seak, Owen Lok In Lo, Wade Chun-Wai Suen, Ming-Tsung Wu

## Abstract

Artificial neural network (ANN) is nowadays one of the most used and researched computational methods. In the field of biocomputing, however, synthetic biologists are still using logic gate technologies to design genetic circuits. We here propose and computationally validate a novel method to mimic ANN with genetic circuits. We first describe the flow and the mathematical expression of this genetic circuit design and then we provide *in silico* proof to support the functionality of our method in regression and classification analyses. We believe that this *de novo* genetic circuit design method would have wide applications in biotechnology.

## Introduction

In the last decade, genetic circuits have been used to simulate logic gates (1–3). In synthetic biology, these logic-gate genetic circuits have been applied to many different aspects including advanced cell therapy (4), chemical sensing (5), and clinical diagnosis (6). Although these studies promote the use of biocomputing from theoretical to clinical applications, the application of genetic circuit was not widely adapted because of the limited computing power of logic gate methodologies.

Machine learning is one of the most popular fields of research nowadays. Every year, thousands of papers were published to develop machine learning algorithms and their applications (7). Many of these algorithms, such as recurrent neural network and convolutional neural network, was developed based on artificial neural network (ANN). An ANN consists of multiple layers and each of them contains multiple nodes. These nodes are regarded as “neurons” in the ANN algorithm, which compute an output according to its activation function, weighting, biases and the current input (8).

Despite the fast growth of computing power and the great advances in machine learning, biocomputing in synthetic biology is still applying the out-of-date simple logic gates, an analog to the first generation of computers which can be dated back to the 1930s. In this paper, we describe a novel method to use genetic circuits to simulate ANNs and this method can unleash the potential of biocomputing algorithms to the next level. With the support of our computational models, we suggest that our method is feasible and applicable to solve real-world biocomputing problems.

## Results

### Flow to design genetic circuit mimicking artificial neural network

Artificial neural networks (ANNs) are sometimes regarded as black boxes. These “black boxes” are equivalent to a group of functions that can solve classification and regression problems, but it is difficult to explain its mechanism of working by mathematical approach (9). Each ANN consists of multiple layers and each layer has multiple nodes. Each of these nodes contains a function defined by the predefined parameters (such as activation functions, number of layers, and nodes) and the training of the network. When we carry out functional training to the network, it is essential to optimize the weighting and the biases of each node, in order to shape the function inside the nodes for more accurate predictions. As shown in figure 1A, the inputted data goes through the calculation inside each ANN node following the order of the layers: all calculated results will be passed through from the current layer to the next layer for further calculation inside each node until reaching the last output layer. Therefore, whether an ANN is successful or not relies on the mathematical functions inside each node (Fig. 1B; top), which its quality is determined by the activation function (Sigmoid, Relu, Tanh, etc), and the transfer function (Eq.1).

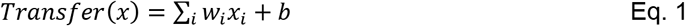

**Figure 1.**
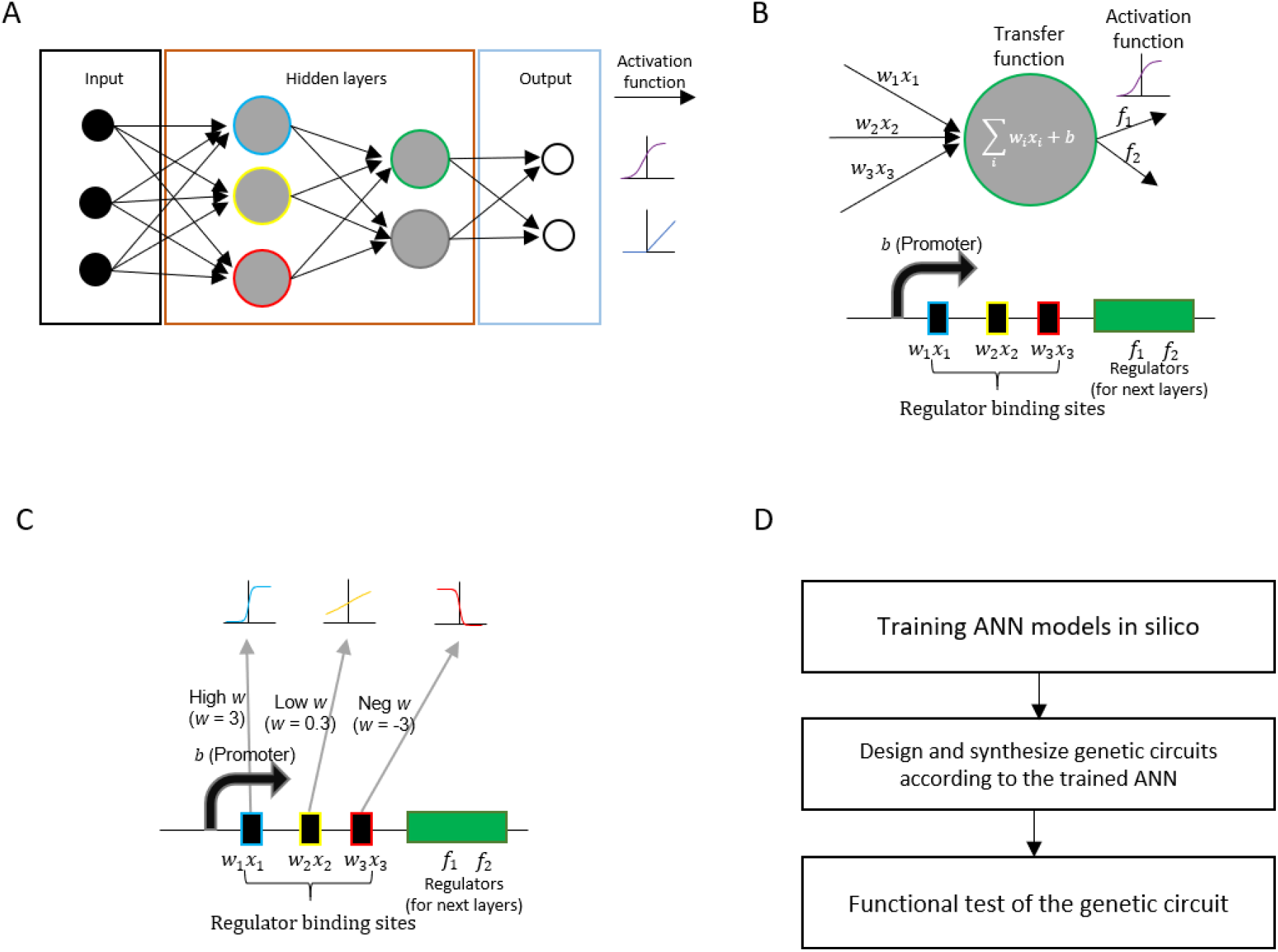
Method to design genetic circuits mimicking artificial neural networks (ANNs). Schematic diagrams showing (A) an example of an ANN *in silico* (B) transfer and activation function in one node of the ANN (top), and the corresponding genetic circuit design to mimic this node (bottom) (C) exemplary regulations in a genetic circuit to mimic the ANN node with different input weightings (see Eq. 3) (D) flow of designing and testing an ANN genetic circuit. (A-C) The colors of the outlines of regulator binding sites in (B-C) represent the respective regulating nodes from the previous layer as shown in (A).

Where in a trained ANN, weighting *w* and biases *b* are fixed constants; *x* represents the input variable from the last hidden layer; *i* represents the number of nodes in the previous hidden layer.

Previous studies (10,11) found that many gene regulations have a shape of sigmoid function (Eq.2), which is also commonly used in ANNs *in silico*. We, therefore, chose the sigmoid function as the activation function to train our ANNs and mimic the transfer and activation functions using transcriptional or translational regulators.

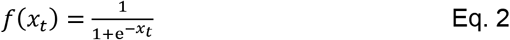

In this sigmoid activation function, the output of the transfer function (Eq. 1) is used as the input *x^t^*. The e represents the *Euler’s* number.

We can therefore mimic the ANN using genetic elements by matching the characteristics of genetic regulators to the trained ANN model, forming a genetic circuit that is the analog to a well-trained ANN in computer (Fig. 1B). The regulation status in each regulator binding site (e.g. transcription factors binding sites; RNA binding sites) can mimic the input of each node, and thereby the number of binding sites will be on the number of nodes in the previous layer. The promoters mimic the biases and the regulators (eg. transcription factors; regulatory RNAs) mimic the activation function.

We can then repeatedly match the reconstructed function from each node to the genetic regulatory elements (Fig. 1C). In practice, because each pair of regulator and binding site only has a single genetic regulation, the mimicking of the transfer function and activation function are conducted within one single function. We choose a regulator and its binding sites based on the weighting function from the current layer and the activation function from the previous layer, and then we get Eq. 3 (hypothesizing the regulations do not interact).

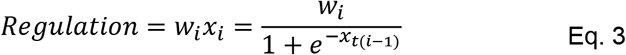

Where *x*_*t*(*i*−1)_ equals to transfer function from the node in the previous layer, which represents the quantity of the regulator from that previous node. When training an ANN *in silico*, *w_i_* is the only variable used to improve the prediction power. Therefore, if we regard *x*_*t*(*i*−1)_ as *x*’, regulation as *y*’ (*y*’ is a function of *x*’), matching the *w_i_*; from pre-trained ANN to genetic regulations (regulators + binding sites) with sigmoid function, we can mimic the ANN using genetic circuits (as shown in Fig. 1C). The progress of matching the genetic regulators can be done by correlation analyses which correlate the functions estimated with Eq. 3 to the regulatory function according to the library of regulators and their binding sites.

Furthermore, we here also propose the flow (Fig. 1D) to generate a new validated ANN genetic circuit and it has three steps: (1) Training ANN models in the computer (2) Design and synthesizing genetic circuits according to the trained ANN (3) Functional testing of the genetic circuit.

### Computational validation of ANN genetic circuit

We also built an ANN *in silico* to prove the feasibility of our proposed method. First, we simulated a data set consisted of three type of inputs (*x*_1_, *x*_2_, *x*_3_) and one output (*Y*) in python. A set of 500 triplet values(*x*_1_, *x*_2_, *x*_3_), were pseudorandomly picked from any real number between 0 to 1. The actual Y value was calculated as:

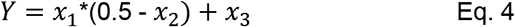

We chose this equation (Eq. 4) to prove that ANN genetic circuit is applicable to perform complicated calculations. We then used these simulated data to train ANN networks with Keras in python (Fig. 2A; two hidden layers and each contains four nodes; epochs=1000) with the leave-one-out method as the validation approach and mean square error (MSE) as the performance function. To mimic the environment of the genetic regulators, we used the sigmoid function as an activation function in all nodes in the ANN. We also introduced a 10% Gaussian noise to each output in the neural nodes to simulate the noise generated by the genetic regulators. We found that even further limiting the activation function by using the “sigmoid” function and adding 10% Gaussian noise, our ANN is still able to predict the simulated data as shown in figure 2B (R=0.9546, p=3.512e-264).

**Figure 2.**
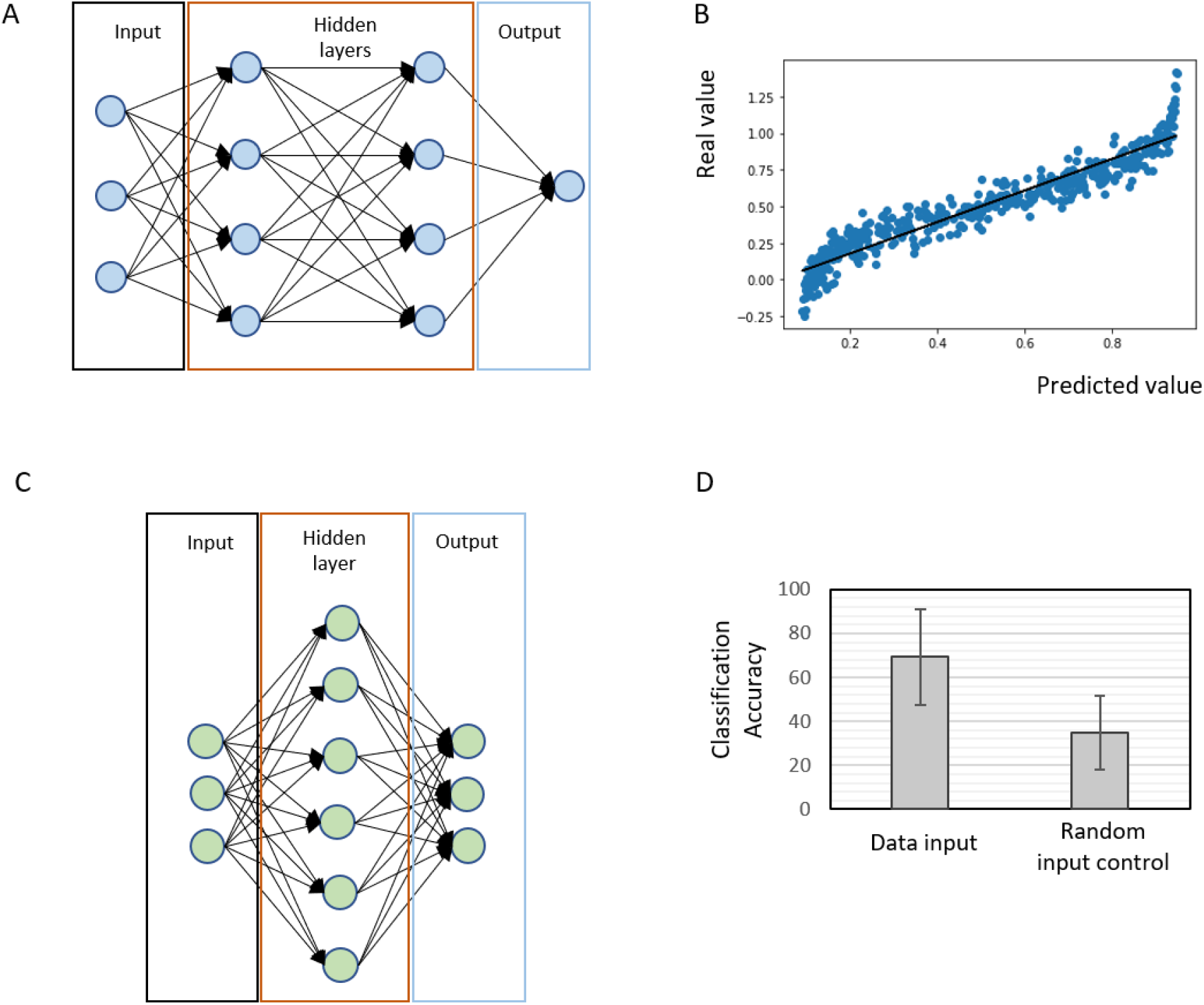
Computational models show that our ANN genetic circuit design can be used for (A-B) Regression analyses and (C-D) Classification analyses. (A) and (C) showing the respective ANN design while (B) and (D) showing the respective modeling result of the two types of machine learning analyses. The black line in (B) represents the Least Squares Regression line. Error bars in (D) represent the Standard Deviations.

To show that our ANN genetic circuit design is a valid solution for real research questions, we used a previously published (12) gene expression dataset of three classified subtypes of breast cancer, basal-like, HER2-enriched, and Luminal (A/B) and selected the expression level of ESR1, PGR, and ERBB2 as input. We built simple ANN circuits with Keras in python (Fig. 2C; one hidden layer which contains six notes; epochs=200) to predict breast cancer subtypes. Like the ANN circuit shown in fig. 2A, we only used the “sigmoid” functions as activation functions in all nodes. We also introduced a 10% Gaussian noise to each output to mimic the noise of the gene regulations. As shown in Figure 2D, with the validated data from the published expression profile, our ANN model performed significantly better in predicting the class of breast cancers (69.29% vs 33.57%; p=0.00165; 10-fold validation) than using pseudorandomly paired expression profile.

## Discussion

In the past decades, the field of biocomputing has been limited to logic gates only, despite the skyrocketing development in computer science. We here described and computationally proved a *de novo* method to mimic artificial neural networks with genetic elements. We used two ANN models to prove the applications of our method in regression and classification analyses and the possibility to solve artificial and real research problems.

It is not the first time that a neural network DNA circuit is being proposed. However, previous researches (13) only used genetic regulators as Boolean classifiers instead of mimicking the ANN with genetic elements. Without the properties we proposed and proved in this paper, the applications of ANN genetic circuits are unfortunately restricted. For instance, if we use the Boolean neural network to design a gene circuit that is similar to the one shown in figure 2C, we will not able to quantify the input. Moreover, such gene circuit will not be able to do any regression (quantification) analysis, which is demonstrated in the ANN shown in figure 2A. Since ANNs designed with our novel gene circuit method can do quantification analyses and perform complicated calculation with fewer neural nodes, we believe our method has a great potential to be applied in many fields in synthetic biology, such as for pharmaceuticals to test drug efficacy, researchers to predict cytokine storms, etc. Furthermore, this biocomputing design can be easily upgraded to other more advanced ANNs, such as recurrent neural networks (RNN; by introducing a regulator binding site that allows the regulator to regulate itself), which gives researchers a great flexibility to adapt the ANN to their researches.

## Author Contributions

L.C.U.S. conceptualized research, L.C.U.S., L.I.L., W.C.W.S., and M.T.W. designed research, performed research. L.C.U.S. wrote the paper. W.C.W.S., and M.T.W. edited the paper.

## Acknowledgments

We thank Prof. Ting Fung Chan, Dr. Jacky Fong Chuen Loo, and Dr. Yen-Po Chin for their comments.

